# Longitudinal whole-exome sequencing of cell-free DNA unravels the metastatic evolutionary dynamics of *BRCA2*-mutated breast cancer

**DOI:** 10.1101/2020.01.10.901330

**Authors:** R.K Hastings, M.R Openshaw, M Vazquez, AB Moreno-Cardenas, D Fernandez-Garcia, L Martinson, B Toghill, K Kulbicki, L Primrose, D.S Guttery, K Page, C Richards, A Thomas, J Tabernero, R.C Coombes, S Ahmed, R.A Toledo, J.A Shaw

## Abstract

Little is known about the metastatic evolutionary dynamics of *BRCA2*-mutated cancers. Here, we applied whole-exome sequencing (WES) of primary tumor (PT), local relapse (LR) and eight serial plasma cfDNA samples collected from disease progression to depict the 12 years evolutionary trajectory of a metastatic *BRCA2*-mutated breast cancer. While longitudinal WES-cfDNA recapitulated clonal and subclonal mutations and copy number profiles detected in LR, emergence of plasma-exclusive mutations in *TSC2* and *HDAC9* cancer-related genes and loss of HLA loci as an immune escape mechanism were also detected. Lastly, mutation signature 3, associated with homologous recombination deficiency and response to platinum-based therapy raised profoundly from 19% in PT to 60% in late stage disease. In conclusion, we show for the first time that longitudinal WES-cfDNA enables the evolutionary trajectory of advanced cancer to be uncovered and that increment of MS3 and loss of HLA are key players in this *BRCA2*-mutated breast metastasis.

## Introduction

Breast cancer is the most commonly diagnosed cancer in women and a leading cause of death. Although advances in treatment and detection have improved survival rates, up to 30% of women worldwide who develop breast cancer die from metastatic disease every year ^1^. Distant relapse can occur up-to 20 years after primary diagnosis and treatment, and it is ultimately responsible for the patient’s death. Once the cancer has returned, it is not always possible to biopsy the recurrence meaning any targeted treatment usually relies on information from the primary tumor, despite the fact that the metastatic cancer may have evolved over time ^2,3^. Analysis of circulating tumor DNA (ctDNA) through “liquid biopsy’ provides an opportunity for non-invasive tumor monitoring, identification of mechanisms of resistance to therapies and detection of actionable mutations that emerge on disease progression ^4-6^.

The development of targeted next generation sequencing (tNGS) enables detection of low frequency alterations at ≤0.1% variant allele fraction (VAF) in ctDNA ^7-9^. Using tNGS, ctDNA has been detected in both early and late stage cancers, including breast cancer ^10-14^ and in metastatic breast cancer (MBC) ctDNA dynamics have been shown to reflect disease progression ^12^. In primary breast cancer, ctDNA profiling has be used to detect molecular relapse, ahead of scans following surgery and/or adjuvant therapy ^4-6^. However due to their limited coverage of the genome, targeted sequencing approaches and patient specific mutation panels may miss key genetic events acquired as the tumor evolves, including potential therapeutic markers.

Aiming to overcome these limitations, new liquid biopsy approaches such as total cfDNA whole-exome sequencing (WES-cfDNA) have been recently developed and shown high concordance with tumor whole-exome sequencing ^15-17^. This WES-cfDNA is able to non-invasively depict the genomic landscape of advanced cancer, monitor known circulating tumor mutations and pinpoint acquired somatic mutations at the time of cancer progression possibly associated with cancer therapy resistance.

We here analysed WES data of 10 DNA samples collected over a 12-year period that comprised the primary tumor (PT, year 2005), local relapse ten years later (LR, year 2015) and eight serial plasma samples collected after relapse, until the patient’s death in 2017. We show for the first time that longitudinal WES-cfDNA enables the evolutionary trajectory of advanced cancer to be uncovered. Moreover, our results also demonstrate that WES-cfDNA can identify tumor mutation signatures, and immune escape mechanism by loss of HLA loci in late-stage cancer. Our study provides insights on the genetic evolution of advanced *BRCA2*-mutated breast cancer and show a prominent role of the *BRCA*-related mutation signature MS-3 and immune escape in the development and evolution of *BRCA2*-mutated breast metastasis.

## Results

This WES investigation concerns a 52 year old female breast cancer patient who harbored a *BRCA2* germline mutation (deletion of Exons 14-16). She was first diagnosed in June 2005 with a 25mm, grade 3, ER positive (5+2), HER2 3+, node positive lobular breast cancer. Following surgery for her primary cancer she was treated with adjuvant cyclophosphamide, doxorubicin and 5-FU plus trastuzumab followed by 5-years adjuvant tamoxifen. In January 2015 (approximately 10 years later) she relapsed with bone metastasis. A relapse biopsy showed ER positive 8/8, HER2 2+ by IHC FISH negative, breast carcinoma. She was followed up with 8 serial blood samples over a 2-year period on treatment until her death.

### WES sequencing metrics

Using an average of only 15 ng cfDNA, we were able to generate genomic libraries for all eight plasma cfDNA samples, which were collected at stable disease or therapeutic response (samples B1, B2, B5, B7), and disease progression (samples B3, B4, B6, B8). The FFPE primary and relapse tumor tissues and germ line DNA isolated form buffy coat also generated successful genomic libraries using 20ng DNA. All sequenced samples passed QC metrics (Supplementary Tables 1 and 2) and we achieved a mean on-target coverage of 107X (range 56-137) per sample.

### WES reveals genomic profiles of tumor and plasma cfDNA

Applying WES-cfDNA to serial plasma samples enabled us to relate genomic profiles to the patient’s clinical disease course (Figure 1). The first CT showed lung progression. Moreover, by the time of plasma B3, taken several weeks before this scan, multiple mutations were detected in ctDNA and a further CT scan soon after showed definitive disease progression within the lung and appearance of a right sided pleural effusion. Following this, the patient was given a course of TDM1, which was clinically ineffective as the subsequent CT showed progressive disease prior to B5. Following a further switch to carboplatin therapy a significant response of the lung disease was seen on CT, reflected by reduced mutations being detected in B5. Plasma B6 showed an increase in the number of mutations and the patient then progressed again with a CT scan showing lesions in her lung and brain and she was switched to letrazole therapy. The plasma cfDNA following a switch to letrazole (B7) showed a reduction in the number of mutations detected suggesting she was responding at that time. However, 10 weeks later a further CT scan confirmed disease progression to brain, which was reflected by a very large increase in mutations in the subsequent plasma (B8) (Figure 1A-C).

**Figure 1.**
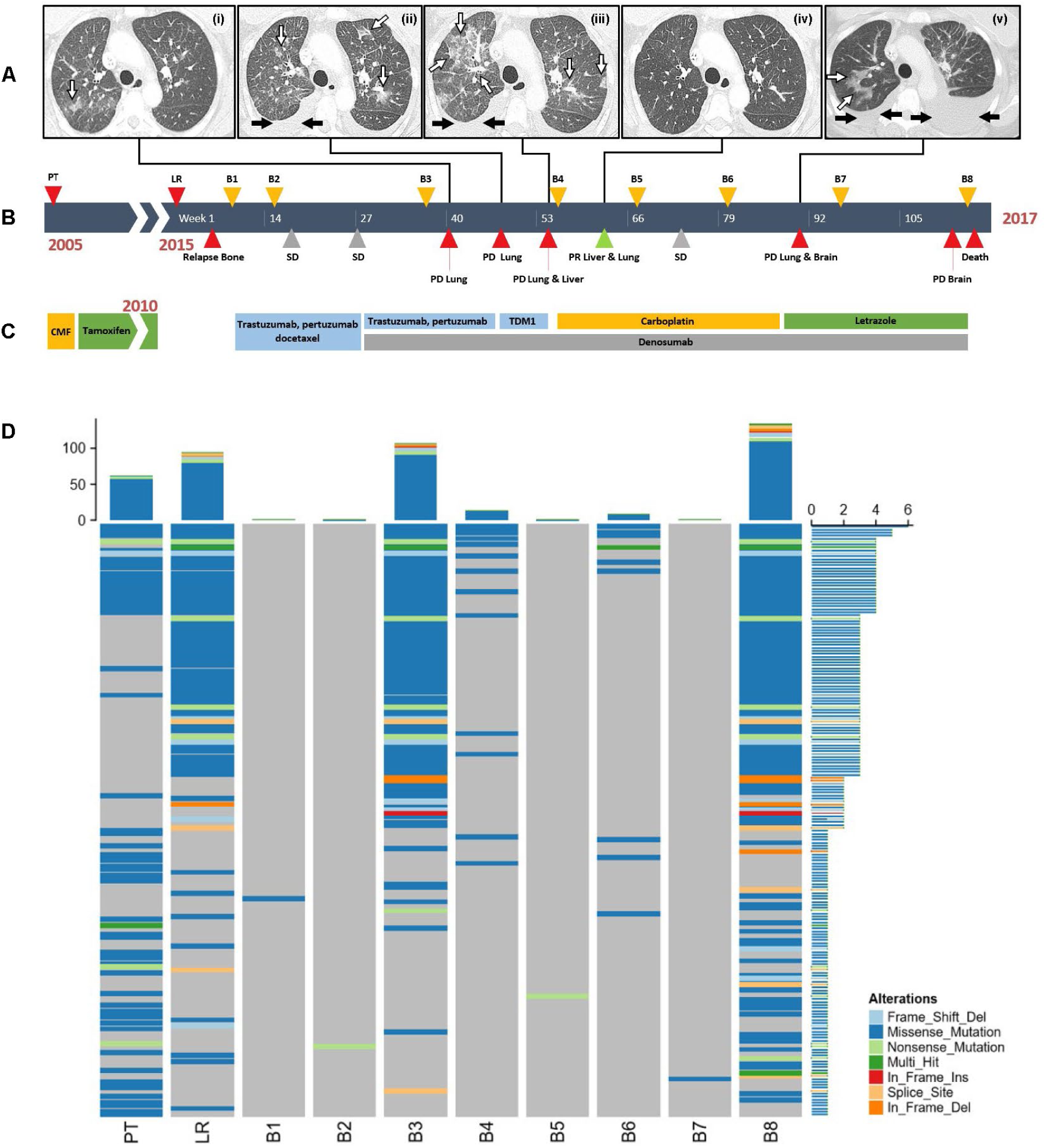
WES captures the genomic landscape of tumor and serial plasma cfDNA. (A) Sequential CT scans (left to right) showing: (I) disease progression potentially lymphangitic carcinomatosis or an area of infection in the lung (arrowed white); (II) areas of lymphangitic carcinomatosis within the lung (arrowed white) and pleural effusion (arrowed black); (III) progressive disease with extensive lung lymphangitic carcinomatosis (arrowed white) and pleural effusion (arrowed black); (IV) resolution of lymphangitic carcinomatosis, and pleural effusion following carboplatin therapy; (V) progression of lymphangitic carcinomatosis (arrowed white) and appearance of bilateral pleural effusions (arrowed black) (B) Clinical history of a female breast cancer patient harbouring a BRCA2 germline mutation from diagnosis in 2005 to her death in 2017 showing dates of blood samples and progression (C) Treatment timeline, (D) Summary of alterations (SNVs and INDELs) detected in tissue and plasma, after applying filtering criteria.

A total of 667 Somatic SNVs and indels and 395 Somatic CNAs (SCNAs) were detected across the 10 samples (PT, LR, B1-B8) (Figure 1D, Table 1, Supplementary Tables 3-6 and Supplementary Figure 1). Applying stringent filtering criteria of having ≥50 SNVs detected, four samples, PT, LR and plasmas B3 and B8 both taken at the time of disease progression were considered for further analysis (Figure 1D). Across these 4 samples the median VAF was 12-36% (range 5-81%, Table 1). Correlation analysis of somatic mutations (SNVs and indels) was performed taking into consideration the VAF of each variant (Figure 2A). All samples were highly correlated: B3 and B8, collected at the time of two different progression disease moments showed the highest correlation (r^2^=0.68 P <0.001), followed by B3 and LR (r^2^=0.63, P <0.001), and then LR and B8 (r^2^=0.49, P <0.001).

**Table 1.** Summary of variants and driver mutations detected by WES. Total number of SNVs and Indels, detected in each sample by WES analysis, showing the median variant allele fraction (VAF) and range (% of total reads), T1 and T2 drivers lists the Tier 1 and Tier 2 driver genes identified.

| **Sample** | **Total variants** | **Total SNV** | **Total Indel** | **Median VAF** | **VAF range min** | **VAF range max** | **T1 Drivers** | **T2 Drivers** |
| --- | --- | --- | --- | --- | --- | --- | --- | --- |
| **PT** | 121 | 119 | 2 | 12.12 | 5.19 | 51.35 | *PIK3CA*<br>p.E545K |  |
| **LR** | 139 | 130 | 9 | 15.58 | 5.13 | 33.77 | *PIK3CA*<br>p.E545K | *DNM2* p.G146D<br>*DCC* p.E495K<br>*ATP1A1*<br>p.W105X *FAT1*<br>p.A173T |
| **B1** | 1 | 1 | 0 | - | - | 10 |  |  |
| **B2** | 2 | 2 | 0 | 7.69 | 6 | 7.69 |  |  |
| **B3** | 155 | 146 | 9 | 34.53 | 5.05 | 81.36 | *PIK3CA*<br>p.E545K | *DNM2* p.G146D<br>*DCC* p.E495K<br>*ATP1A1*<br>p.W105X *TSC2*<br>p.V461M |
| **B4** | 17 | 17 | 0 | 6.35 | 5.05 | 11.11 |  |  |
| **B5** | 2 | 2 | 0 | 8.33 | 6.67 | 8.33 |  |  |
| **B6** | 14 | 14 | 0 | 6.36 | 5.41 | 10.29 | *PIK3CA*<br>p.E545K |  |
| **B7** | 3 | 3 | 0 | 9.31 | 7.14 | 10.29 |  |  |
| **B8** | 213 | 193 | 20 | 19.58 | 5.05 | 52 | *PIK3CA*<br>p.E545K | *DNM2* p.G146D<br>*DCC* p.E495K<br>*ATP1A1*<br>p.W105X *HDAC9*<br>p.S525C |

**Figure 2.**
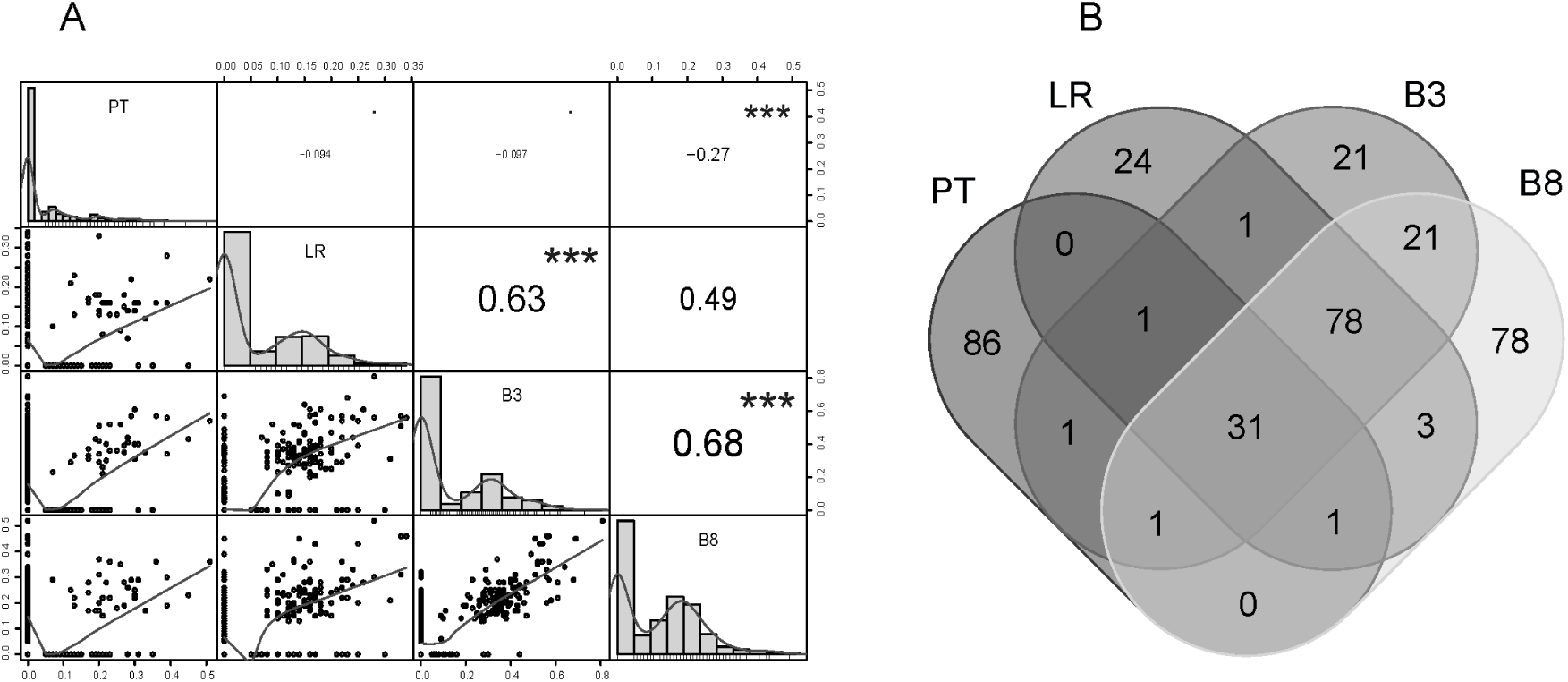
High correlation between serial plasma and tumor at relapse. (A) Correlation analysis of somatic mutations (SNVs and indels) including consideration the VAF of each variant. (B) Overlap of mutations present in PT, LR, and plasma B3 and B8 samples.

We compared the overlap of variants from each of the four genomes. Of 121 mutations identified in the PT, only 33 (27.3%) were detected in the LR diagnosed 10 years later and 34 (28.1%) and 33 (27.3%) overlapped with B3 and B8 respectively (Figure 2B). WES-cfDNA of B3 and B8 were highly informative and captured 79.9% (111/139) and 81.3% (113/139), respectively, of the mutations detected in the LR. Moreover, WES-cfDNA uncovered highly confident plasma-exclusive mutations: 42 in B3 and 99 in B8, 21 of which were common to both (Figure 2B). These longitudinal results suggest the occurrence of a small core group of mutations that persist from first diagnosis to the patient’s death 14 years later, and also reveal the dynamics of acquired sub-clonal mutations in *BRCA2*-mutated breast disease.

### WES-cfDNA empowers tumor evolution analysis

To investigate the degree of tumor evolution, 321 eligible SNVs that were present in at least one of the 4-time points (PT, LR and plasmas B3 and B8) along with their corresponding VAFs were analysed using LICHEE ^18^ (Figure 3A). Thirty SNVs were detected to be common to all 4-time points with the *PIK3CA* p.E545K driver mutation present in the first cluster. This cluster persisted through all time points despite switches in treatment. We observed 85 SNVs that were unique to the PT (i.e. variants within the cluster that are not shared with any other sample time point) and no Tier 1 or Tier 2 drivers were detected within this cluster. Seventy-six additional SNVs were present in LR, B3 and B8 time points and this cluster contained the TIER 2 driver genes *DCC, DNM2, ATP1A1*. Twenty SNVs were detected only at LR and the Tier 2 driver *FAT1* was found only at this time point. We observed 16 additional variants in common with B3 and B8 not present in PT and LR. B3 had 20 SNVs not detected at any other time-point, including a Tier 2 driver *TSC2* variant. B8 had 68 SNVs with a tier 2 driver *HDAC9* variant detected only at this time-point. The predicted tumor lineage reflects the known sample time points and provides evidence of tumor evolution, aligning with clinical progression of the patient.

**Figure 3.**
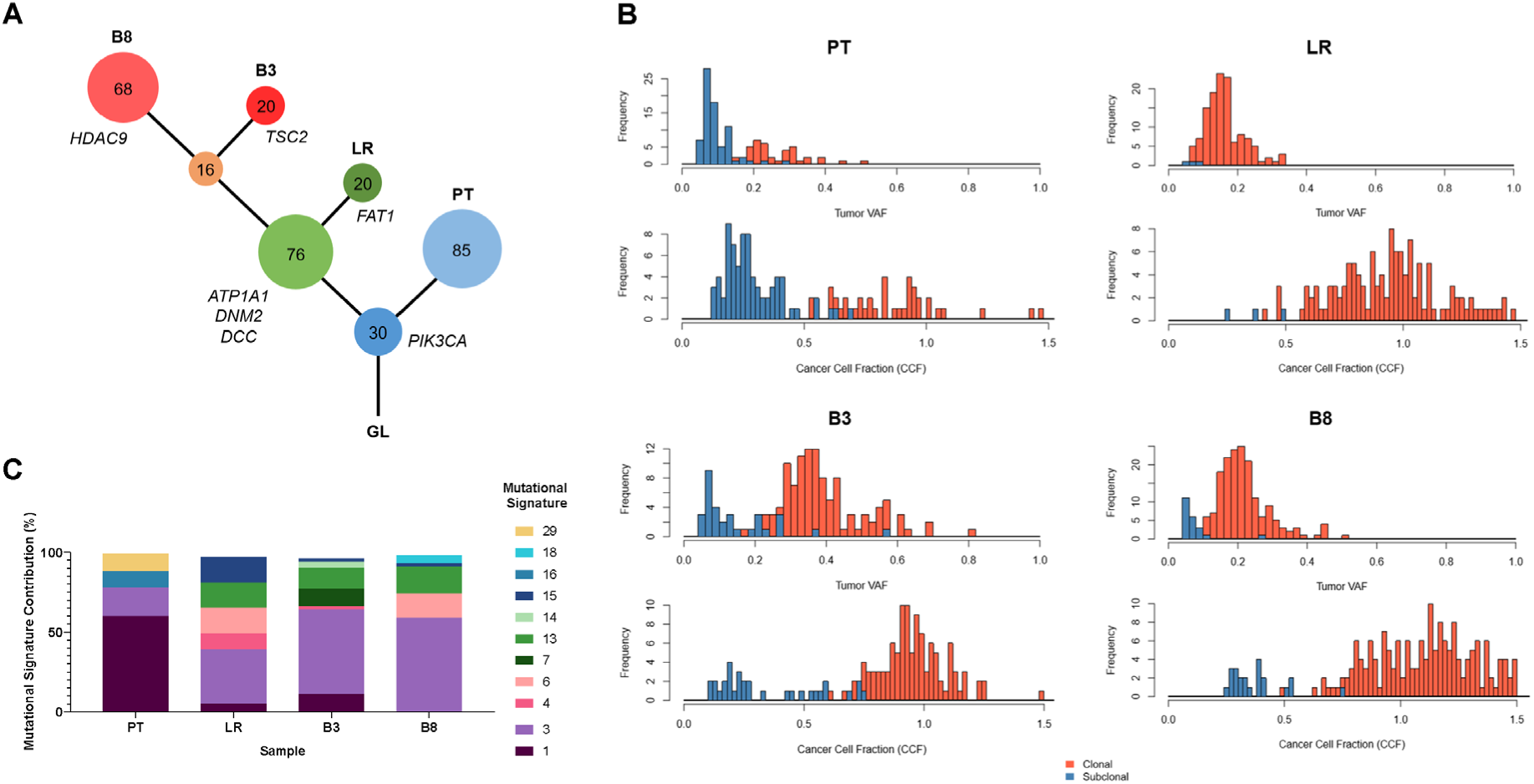
Tumor evolution is characterised by emergence of driver mutations and clonal mutational signatures at disease relapse. (A) Phylogenetic tree of the WES samples, noting emergence of driver mutations with disease progression. Large circles link the clonal changes that are acquired on progression, smaller circles show private mutations; (B) The cancer cell fraction (CCF) distribution shows a clear trend of progression from predominantly sub-clonal variants in PT to predominantly clonal variants with disease progression in LR, B3 and B8; (C) Mutational signature analysis revealed that MS-1 was the predominant signature in the primary tumor at 60.17% but this resolved by B8. MS-3, which is associated with failure of DNA double-strand break-repair by homologous recombination was also detected in PT (18.64%) and increased to 59.59% becoming the major signature in B8 taken shortly before the patient died.

### Emergence of cancer driver mutations and mutational signatures at disease relapse

Somatic mutations from PT, LR, and plasmas B3 and B8 were analysed using the Cancer Genome Interpreter (CGI) ^19^ to detect known and predicted cancer driver genes. A single Tier 1 driver mutation, *PIK3CA* p.E545K, was detected in all 4 samples at a VAF of 21.6% - 53.8% (Supplementary Table 5). This mutation was also identified at 7.7% in B6 that had only 17 somatic mutations detected by WES-cfDNA. The *PIK3CA* p.E545K mutation was confirmed by targeted NGS (tNGS) using an Ampliseq panel ^20^ (Supplementary Table 7), which also identified sub-clonal *ESR1* gene mutations,subsequently confirmed, by ddPCR (Supplementary Table 8).

Three Tier 2 driver mutations *(DNM2 p.G146D, DCC p.E495K and ATP1A1 p.W105X*) were detected from WES (Supplementary Table 5) and stratified as clonal variants (Figure 3A). Interestingly, all three mutations were found in LR, plasma B3 and B8 but were undetected in the PT, suggesting that they emerged during the evolution of the disease. Other Tier 2 mutations were *FAT1 p.A173T* that was detected exclusively in the LR, *TSC2 p.V461M* detected exclusively in B3, and *HDAC9 p.S525C* that emerged exclusively in B8 (Figure 3A).

Of interest, the cancer cell fraction (CCF) distribution shows a clear trend of progression from predominantly sub-clonal variants in the PT to a majority of clonal variants with disease progression in LR, B3 and B8 (Figure 3B). In addition, mutational signature analysis revealed that MS-1 was the predominant signature in the primary tumor at 60.17% (Supplementary Table 9 and Figure 3C) comprising both clonal (n=23) and sub-clonal mutations (n=48) (Supplementary Figure 2 and Supplementary Table 10). MS-1 is reported to be associated with an endogenous mutational process initiated by spontaneous deamination of 5-methylcytosine ^21^. MS-3, which is common to *BRCA*-mutated cancers and is associated with failure of DNA double-strand break-repair by homologous recombination, was also detected (18.64%) comprising clonal (n=10) and subclonal mutations (n=12). Other prevalent signatures in the PT were MS-16 (10.17%), and MS-29 (11.02%) (Supplementary Table 10).

The local relapse (LR) showed changes to the mutational signatures, with MS-1 reduced (5.43%) and MS-3 predominating (34.88%). Emergent signatures at LR included MS-13 (16.28%) found in many cancers ^21^ and attributed to APOBEC activity, MS-15 (16.28%) associated with defective DNA mismatch repair, MS-6 (16.28%) associated with defective DNA mismatch repair also known to be present in samples with MS-15 ^21^, and MS-4 (10.85%) associated with exposure to tobacco carcinogens. At LR, the number of sub-clonal mutations was minimal with only 3 predicted out of 129 mutations. In plasma B3, taken 5 months after disease progression, MS-3 predominated (53.47%), whilst MS-1 (11.81%) increased and MS-15 (2.78%), MS-4 (2.08%) and MS-13 (13.89%) decreased. Emergent new signatures at B3 were MS-7 (11.11%) and MS-14 (4.86%). MS-6, detected at 16.28% in the LR was not detected. The number of sub clonal events for MS-3 increased to 25.97% (20/77) (Supplementary Figure 2 and Supplementary Table 11).

Our data show major changes in the mutation signature landscape from PT to subsequent relapse and late stage disease progression in *BRCA2*-mutated breast cancer. MS-3 rose from 18.64% in PT to 59.59% and was predominately clonal in contribution, becoming the major signature in plasma sample B8 taken shortly before the patient died. MS-13 increased (17.62%) and MS-6 re-emerged (15.54%). MS-15 was greatly reduced (2.07%), MS-18, reported previously in breast and stomach carcinomas emerged (5.18%) and MS-1, which was the dominant signature in the PT was not detected (Figure 3C).

### Neo-epitope analysis

The germ line DNA sample was used to determine the alleles of HLA genes encoding the MHC Class I complexes. The alleles predicted for MHC Class I HLA genes, specified up to the second field were HLA-A*02:01, HLA-B*14:01, HLA-B*08:01, HLA-C*08:02, and HLA-C*07:01. The HLA-A gene was predicted to be homozygous, whilst HLA-B and HLA-C were predicted to be heterozygous.

We considered 628 variants (347 unique) in the samples under study: PT, LR, B3, and B8. From these, 216 different mutations produce potential MHC Class I binding peptides. The scores for MHCflurry, which were successfully computed for just 187 of the 216 peptides (all peptides are successfully predicted by at least one method), are shown in Supplementary Figure 3A. Of the 216 mutations, 93 (43%) co-occur on several samples while the rest are unique to one sample; 74 (34%) were exclusively found in plasma, while 66 (30%) were only found in PT and LR. The epitopes common to both tumors were also detected in plasma (Supplementary Figure 3B, Supplementary Table 11).

### WES-cfDNA provides an opportunity to detect immune evasion as the tumor progresses

To establish if WES-cfDNA samples could be used to detect immune evasion by the tumor, we used the HLA Class I allele predictions from Polysover ^22^ for the germline sample and plasma time points (Supplementary Table 12). As only HLA-B and HLA-C were heterozygous, we then used the predicted alleles for HLA-B and HLA-C with the predicted ploidy and purity from Sequenza for each tumor and WES-cfDNA timepoint and used the program LOHHLA ^23^ to establish if there had been any loss of heterogeneity at each time point. Evidence of LOH was detected in the HLA-B gene in PT, B3 and B8 samples, although only B8 reached statistical significance (P <0.05) and for HLA-C LOH was detected in all 4 samples (P<0.05) (Supplementary Figure 4, Supplementary Table 12).

### Clonal SCNAs persist with breast cancer progression

Somatic copy number alterations (SCNAs) were identified using the allelic copy number caller Sequenza. A number of clonal SCNAs were detected by global genomic copy number profiling, including to 1q, 3q, 8q and 19q that persisted from PT through to B3 and B8 with breast cancer progression (Figure 4, Supplementary Figure 5).

**Figure 4.**
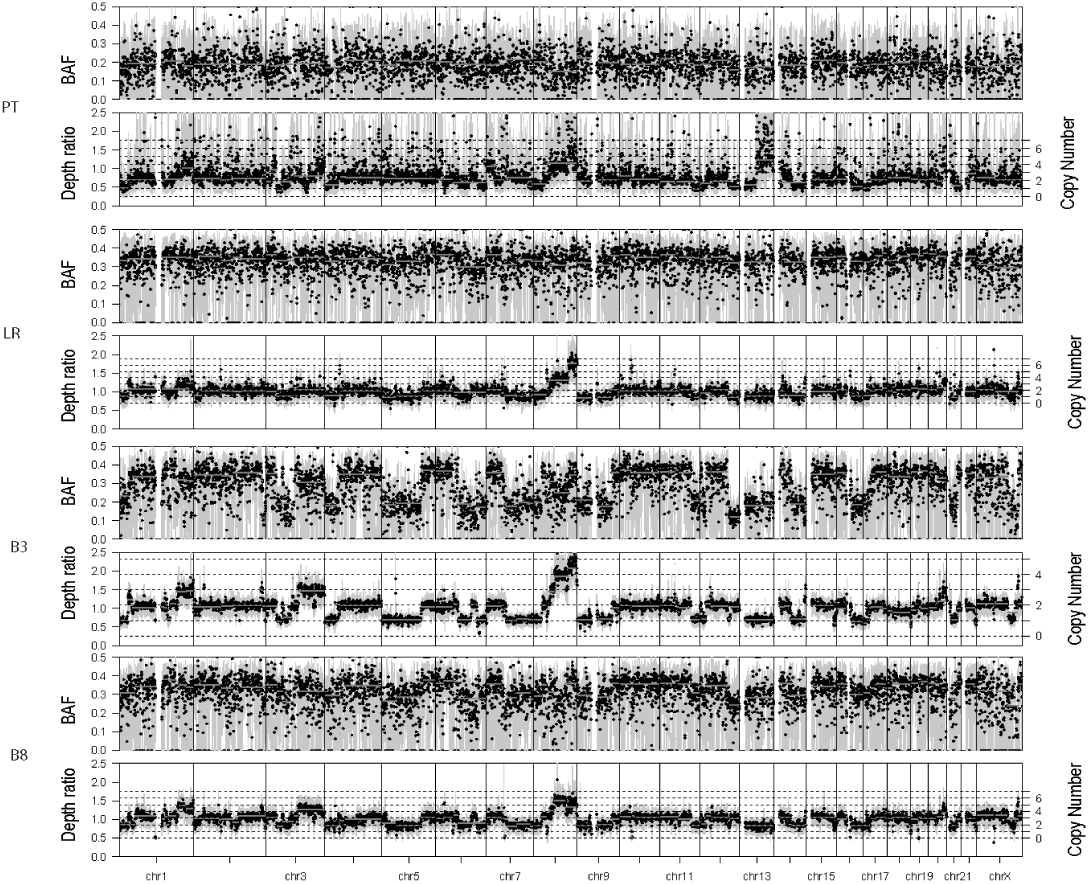
Clonal SCNAs persist with breast cancer progression. Somatic copy number alterations (SCNAs) were identified using the allelic copy number caller Sequenza.

We defined the copy number calls into thresholds; amplifications (defined as CN=2*ploidy + 1), gains (CN + 1 & < CN amplification threshold), loss (CN < ploidy & >0) and deletions (CN=0) (Supplementary Table 6). We also defined the copy number calls into focal (<=25% of the chromosome arm size) and broad regions (>=25% of the chromosome arm). Calls less than 10kb in length were excluded. We detected 395 regions of SCNAs across the 10 samples with a mean of 39.5 per sample (median=35.5). We observed 72 amplifications (71 Focal and 1 Broad), 123 gains (113 focal, 10 broad), 189 losses (96 focal, 93 broad) and 11 homozygous deletions all focal, however there were no broad deletions (Supplementary Table 6 and Supplementary Figure 1).

We analysed the genes in the copy number altered regions from the exome analysis in the CGI to establish any known SCNA drivers. We observed no deleted genes indicated to be drivers. A focal amplification containing the pro-oncogene *MYC* was detected in the LR at CN 6, and a focal gain in plasmas B3 at CN 4 and B8 at CN 4. We also detected *MYC* in the PT at a CN 4. The highest level of amplification detected across all 4 samples (CN of 4-16), spanned 8q24.3 (chr8:145665359-145993019) and contains 13 genes, none of which are known cancer related genes. (Figure 4, Supplementary Table 6).

The region 3q13.13-q29 (Chr3:109052732-1097847031) containing 728 genes showed a gain in the PT, B3 and B8 with a CN of 3-4 and contains the *PIK3CA* and *SOX2* genes. Region 1q41-q42.11 (chr1:229366646-224202533) was amplified in samples LR, B3 and B8 with a CN of 7-8. Three genes were present in this region, two pseudogenes and TP53 binding protein *TP53BP2*.

There was uncertainty concerning the HER2 status of the patient at relapse. The FFPE primary tumor was reported as HER2 positive (HER2 3+ by immunohistochemistry analysis), whereas the relapse biopsy was reported as HER2 2+ and FISH negative. Review of the WES SCNA calls showed no amplification in the region encoding the *ERBB2* gene (17q12, chr17:37856254 – 37884915) at any time point (Figure 4), suggesting a HER2 negative status throughout her disease course. Clinically, a HER2 negative status for the patient is in keeping with the lack of clinical response seen to trastuzumab/pertuzumab and TDM-1 (Figure 1, time points B3 and B4).

## Discussion

The usefulness of plasma ctDNA analyses using a targeted ddPCR or gene panel strategy are well established ^4-14^ but our study highlights the potential advantage of longitudinal WES-cfDNA as exemplified by this patient with metastatic breast cancer. WES-cfDNA combines the convenience of less-invasive plasma cfDNA studies with the robustness of an exome sequencing approach. We also show that WES-cfDNA genomic profiling is capable of identifying acquired mutations during breast cancer progression, as well as providing information as to clonal evolution, tumor mutation signature analysis, identification of neo-epitopes and detection of immune escape mechanisms.

In addition to recent technical advances ^17^, our longitudinal WES-cfDNA approach generated biological insights of the patient’s disease course from relapse until near to the time of her death. We found a core of 31 mutations and specific SCNA profiles that remained throughout her disease development from the PT, LR and through treatment for metastatic disease, comprising a period of 14 years. Strikingly, this clonal driver core represented only 8.6% (31 of 361) of all the *bona fide* mutations identified within this period. This suggests that the history of this patient’s *BRCA2*-mutated breast cancer is mainly driven by an enduring clone with the acquisition of many sub-clonal mutations that fluctuated during the course of her disease and treatment. In this regard, drugs against these core molecular targets would have had potentially high therapeutic impact controlling tumor growth.

WES-cfDNA allowed the identification of Tier 1 drivers that were present at initial primary tumor biopsy and additional Tier 2 drivers emerging at relapse and as the cancer progresses. We identified a clonal Tier 1 driver in the *PIK3CA* gene p.E545K present at diagnosis in 2005, persisting when the cancer returned some 10 years later in the LR, and in the WES-cfDNA samples B3 and B8 despite switches in treatment. As a clonal driver, *PIK3CA* p.E545K was also observed at time point B6, which had only 17 somatic mutations detected by WES-cfDNA. Putative pathogenic mutations were detected in *DNM2 (p.G146D), DCC (p.E495K) and ATP1A1 (p.W105X)* in the LR and B3 and B8 plasma samples but were not present in the PT, which suggests evidence of tumor evolution and an implication with cancer relapse after many years dormant. The *DNM2* gene has been linked with cell migration and metastasis, receptor endocytosis and with resistance to endocrine therapy receptor endocytosis ^24,25^. The *DCC* gene is frequently deleted or its expression reduced in breast cancer ^26^. *ATP1A1* genes has been shown to have decreased expression in human renal cell carcinomas compared to the adjacent non-tumor tissues and it was therefore proposed that *ATP1A1* is a potential novel suppressor protein for renal cancer ^27^. One Tier 1 driver mutation in *FAT1 (p.A173T)*, was exclusive to the LR, and not detected in the subsequent serial plasma samples. This mutation, was not detected in the PT and appears to be a sub-clonal mutation that resolved whilst the patient was on Pertuzumab, trastuzumab and docetaxel at time point B3. *PIK3CA* and *FAT1* mutations are frequent in hormone receptor positive patients ^28^, as in this case, where the PT was reported as ER+ (5+2). We also observed a mutation in *TSC2* at B3 only, just prior to disease progression to her lung. *TSC2* acts with *TSC1* in a complex to inhibit mTOR, an emerging therapeutic target and known promoter of cell growth and cell cycle progression ^29^, This suggests that the patient would have benefited from everolimus. At timepoint B8 when the patient was on Letrozole therapy, we also observed a mutation in the *HDAC9* gene. Increased expression of the *HDAC9* gene is associated with antiestrogen resistance of breast cancers ^30^, consistent with the late acquisition of low frequency sub-clonal mutations in *ESR1*, detected by ddPCR in B8. Knowledge of the key driver mutation in *PIK3CA* p.E545K, could have provided information for therapy selection. For example a recent ctDNA based study suggested that the mTOR inhibitor everolimus, may be a useful chemotherapy adjunct ^31^ which suppresses *PIK3CA, ESR1*, and *GATA3* gene mutations and may have been a viable alternative therapy. Alternatively, treatment with the newly developed PI3K inhibitor drug alpelisib ^32^ could have been indicated alongside fulvestrant.

Sequential WES-cfDNA also allowed the characterization of dynamic changes in mutational signatures throughout the disease progression, including 2 plasma samples that had sufficient variants detected in ctDNA (B3 and B8). Contrary to what we expected for a breast tumor from a patient carrying a *BRCA2* exon 14-16 germline deletion, the MS-3 signature, associated with the failure of DNA double stranded break-repair by homologous recombination ^21^, was underrepresented in the PT (18.64%) and only became dominant later on disease progression, increasing from 34.88% at LR, to 59.59% at B8. After progression to lung and liver on TDM-1 (timepoint B3), the patient was given carboplatin, a chemotherapy specifically shown to be active in patients with MS-3 BRCA-related disease ^33^ and she initially responded well to this therapy as determined by imaging and reflected in ctDNA results. However, overall no given treatment eradicated the MS-3 signature suggesting that MS-3 is likely to be reflecting the mutational signature of the lethal clone. Conversely, MS-1, that was the most prominent in PT, decreased with disease progression until it was completely undetectable in sample B8 just prior to the patient’s death. In line with this, it will be interesting in the future to use longitudinal WES-cfDNA to monitor tumor mutation signatures and especially the MS-3 in the context of *BRCA2*-mutated MBC treated with PARP inhibitors. Overall, there were 7 predominantly clonal mutational signatures detected in plasma as the disease evolved not present in PT, and 3 of these were also absent from LR. Identifying additional mutation signatures captures the evolving intrinsic mechanistic processes involved in the cancer progression by identifying mutational signatures that were not present in the PT.

Our data also showed that the sequential WES-cfDNA approach can depict the dynamics of predicted tumor neoepitopes during the cancer evolutionary trajectory. As the genome of the breast cancer evolved and acquired new mutations, neoepitopes are potentially presented by the HLA proteins, recognized and become a focus for tumor infiltrating lymphocytes (TILs) and other cells of the immune system. Our study shows that sequential WES-cfDNA can be a powerful tool for the identification of neoantigens that can be used for the design of cancer vaccines ^34^.. We also show for the first time that WES-cfDNA can be used to monitor dynamic changes in the copy number of HLA alleles, highlighting the opportunity to detect immune evasion as the tumor progresses. Based on these results, we are currently testing WES-cfDNA for monitoring of cancer patients treated with immunotherapy.

Additional benefits highlighted by this study show that genomic copy number profiles can be generated from WES-cfDNA that capture key aberrations, including detection of clonal drivers. For example, amplification was detected in 1q, 3q, 8q and 19q ^2^ that persisted from PT through to LR, B3 and B8. Moreover, WES-cfDNA profiling revealed that this patient was likely to be HER2 negative, but had amplifications and gain alterations in *MYC* and other potentially important genes including *MDM4, SOX2* ^35^ *and AURKA* (Figure 4). The WES-cfDNA *ERBB2* result predicted non-response to TDM-1, an anti-HER2 antibody-chemotherapy conjugate drug indicated for use in HER2 positive breast cancers ^36^ given following sample B3. Since sample B3 also had a strong MS-3 profile, carboplatin could have been used at an earlier time-point instead of TDM-1. Several potential drugs are under development for the treatment of patients with *MYC* amplification including Omomyc ^37^ but none have yet reached clinical use. Acting on such information provided by WES-cfDNA and thereby selecting the right drug at the right time, may have prevented clinical deterioration, and highlights the possibility of developing WES-cfDNA for personalised precision medicine.

A clear limitation of WES-cfDNA is the requirement for sufficient ctDNA levels in order to generate genomic profiles. As shown in our study plasma samples taken at times of disease progression (B3 and B8) were most informative and revealed a number of mutations and amplifications of key driver genes, whereas those plasma samples taken when disease was well controlled had few detectable variants. In these samples additional experiments such as targeted deep sequencing or ddPCR can be considered as used here, where emergent sub-clonal *ESR1* gene mutations were detected in B8 that were not detected by WES-cfDNA.

In summary, our study shows that sequential WES-cfDNA is a reliable technology that can recapitulate the critical genetics of the underlying metastatic disease, and track the evolving changes in the cancer genomic landscape throughout different treatments. Sequential WES-cfDNA may provide data that can help the real-time selection of therapies to tackle the evolving cancer disease of the patient, which as we showed here can evolve and change over a short period of time. Importantly, this study shows that *BRCA*-related tumor mutation signature 3, which became highly prominent, and loss of HLA are potential influential players during the metastatic evolutionary history of BRCA2-mutated breast cancer.

## Methods

### Patient recruitment and sample collection

The patient was recruited with written informed consent to our study at the time of relapse and followed up with 8 serial blood samples over a 2-year period on treatment until her death. The study protocol was approved by the University of Leicester Cancer Research Biobank UHL11274 tissue access committee (REC reference number: 13/EM/0196), DNA from one region of the primary tumor, one region of the recurrent local relapse tumor and 8 serial plasma cfDNA samples were used for WES.

### Whole-exome sequencing

An area of tumor tissue was identified in the diagnostic FFPE block by a consultant histopathologist and sampled using a 1mm TMA needle core to gain high cancer cellularity. Tumor DNA was extracted using the GeneRead™ DNA FFPE kit (Qiagen®). Total cfDNA was isolated from 3ml plasma using the QIAamp® Circulating Nucleic Acid Kit and quantified by Qubit as described previously ^38^. DNA isolated from cells in the buffy coat served as a germline control. Libraries were prepared using 15ng of DNA following the WES workflow using the Illumina HiSeq4000 platform as described in Toledo, Garralda et al 2018 ^17^.

Paired end raw data in FASTQ format were trimmed with trimmomatic (0.36) ^39^ and aligned to the reference human genome (hg19) using Burrows-Wheeler Aligner, BWA (0.7.15) ^40^. The aligned BAM file files were sorted with Samtools 1.3.2 ^41^. Duplicates and base recalibration was performed using Picard tools v2.7.1 (http://broadinstitute.github.io/picard/) (Accessed October 2019) and applied GATK ^42^ base quality score recalibration, indel realignment, duplicate removal, and performed SNP and INDEL discovery using standard hard filtering parameters or variant quality score recalibration according to GATK Best Practices recommendations ^43,44^.

We used the CollectHSmetrics from the Picard suite of tools to calculate the mean coverage across the aligned BAM for each sample and the percentage on target coverage. Values reported are based on the default setting of CollectHSmetrics using a minimum mapping and minimum base quality of 20.

### Somatic SNV and indel Detection

Somatic SNVs and INDELS were called with Mutect2, part of the GATK4 suite of tools ^42^. Additional filtering criteria was used on the VCF output of Mutect2 for confident somatic calls; for SNVs a depth of ≥50 reads with ≥3 mutant reads and a VAF ≥=5%. For INDELS a depth of ≥50 reads with ≥10 mutant reads and VAF ≥5%. The Cancer Cell Fraction (CFF) was calculated by Palimpsest ^45^ using filtered mutations in conjunction with SCNA segments, predicted tumor cellularity and ploidy information derived from Sequenza ^46^.

### Functional annotation

Functional annotation of SNV and indels was performed with Variant Effect Predictor (v86),^47^ via the vcf2maf program (available at https://github.com/mskcc/vcf2maf) (Accessed October 2019), ensuring that each variant was mapped to only one of all possible gene transcripts/isoforms that it might affect. We used MAFtools ^48^ to summarise the variants.

### Driver gene identification

TIER 1 and TIER2 drivers were identified using the cancer genome interpreter ^19^ for SNVs, INDELs, and SCNA using the web-based analysis function at https://www.cancergenomeinterpreter.org/home.

### Mutational Signature Analysis

To identify mutational processes that may be driving the tumor evolution, SNVs were analysed to extract the mutational signatures using the R package Palimpsest ^45^. We then compared the extracted mutational signatures once decomposed to known signatures derived from Alexandrov, Nik-Zainal ^21^, and cosine similarity was calculated to identify best match and the relative contribution in each sample.

### Phylogenetic analyses based on SNV and VAF

We used LICHEE ^18^ to reconstruct multi-sample cell lineage trees and infer the subclonal composition based on the variant allele frequency (VAF) data of filtered SNVs from PT/LR tumor samples and B3 / B8 plasma samples, collected at different disease progressions. We eliminated private clusters that had fewer than 2 SSNVs and only considered mutations with a VAF <75%.

### Copy number Aberration Identification

To determine the somatic Copy Number Aberration profiles of the WES-FFPE tumor and WES-cfDNA samples we applied the allele-specific copy number R package Sequenza v2.1.2 ^46^. The processed final tumor and plasma BAM files with the matched lymphocyte as a germline sample were used as input. We followed the standard workflow for BAM files in the R Vignette (https://cran.r-project.org/web/packages/sequenza/index.html) (Accessed October 2019). We then mapped the segmented regions to a cytoband location along with the genes within using Bedtools ^49^.

### MHC Class I typing

The control sample (GL) was used to determine the alleles of HLA genes encoding the MHC Class I complexes. To this end, the data were processed using three different tools: SOAPHLA (v1.2), OptiType (v1.3.1), and Polysolver (v4).

### Neo-epitope prediction pipeline

The somatic variants from all samples where submitted to the neo-epitope prediction pipeline using PVacSeq (from PVacTools package v1.3.7). First a VCF if composed with the variants and annotated with VEP following the documentation of the PVacSeq tool. The tool was run for the alleles specified above, with peptides of sizes 8, 9 and 10, and the following methods: MHCflurry, MHCnuggetsI, NetMHC, NetMHCcons, NetMHCpan, PickPocket, SMM, and SMMPMBEC. The MHCflurry ^50^ models were installed locally (v2.19.1).

### HLA Loss of Heterozygosity prediction

To establish if there has been tumor HLA allelic loss over the timeline of the disease we used the predicted Polysover HLA alleles for A, B and C in addition to the predicted ploidy and tumor purity from sequenza and ran the LOH HLA copy number prediction pipeline ^23^ in tumor and plasma cfDNA samples. We used default parameters and followed the instructions for use at https://bitbucket.org/mcgranahanlab/lohhla/src/master/ (Accessed October 2019).

### Orthogonal validation of key mutations by tNGS and ddPCR

20ng serial cfDNA samples were sequenced using a custom 23 amplicon Ion AmpliSeq panel covering hotspot regions in *ESR1, PIK3CA, TP53, and ERBB2* as described previously ^20,51^, and using ddPCR assays (*PIK3CA* p.E454 from Bio-Rad) and in house *ESR1* and *ERBB2* assays.

## Acknowledgements

We thank the University Hospitals of Leicester NHS Trust, the Leicester Experimental Cancer Research Centre and the HOPE Clinical Trials unit for supporting patient recruitment and sample collection. This research used the ALICE and SPECTRE High Performance Computing Facilities at the University of Leicester.

## Authors’ contributions

**Conception and design:** RAT, JAS, MRO, RCC

**Development of methodology:** RAT, RKH, MV, ABMC, KP, LP, DG, LM

**Acquisition of data (acquired and managed patients, provided facilities, etc.):** KK, AT, SA, CR, MRO

**Analysis and interpretation of data (e.g., statistical analysis, biostatistics, computational analysis):** RKH, RAT, MV, DFG, BT, LM, JAS

**Writing, review, and/or revision of the manuscript:** RKH, MRO, RAT, JAS, RCC

**Administrative, technical, or material support (i.e., reporting or organizing data, constructing databases):** RKH, JT, DFG, BT, LP

**Study supervision:** RAT, JAS, AT

## Supplementary Tables

Supplementary Table 1: Whole exome sequencing metrics

Supplementary Table 2: Whole exome sequencing on target coverage metrics

Supplementary Table 3: SNV and Indel summary

Supplementary Table 4: SCNA summary

Supplementary Table 5: SNV and INDELS calls and annotations

Supplementary Table 6: SCNA calls and annotations

Supplementary Table 7: Targeted Sequencing Coverage for PIK3CA and ESR1 mutations

Supplementary Table 8: Digital Droplet results for PIK3CA and ESR1 mutations

Supplementary Table 9: Mutational signatures proportions

Supplementary Table 10: Clonal and subclonal mutational signature distribution

Supplementary Table 11: Neoepitope summary

Supplementary Table 12: Loss of Heterozygosity prediction in the HLA-B gene and HLA-C gene

## Supplementary Figures

Supplementary Figure 1. (A) Number of somatic variants by mutation type (B) number of somatic copy number aberrations by call type.

Supplementary Figure 2. Clonal and subclonal distribution of mutational signatures in PT, LR, B3 and B8.

Supplementary Figure 3. (A) Number of potential MHC Class I binding peptides predicted by MHCflurry (B) Co-occurrence of neoepitopes in PT, LR, B3 and B8.

Supplementary Figure 4 (A). HLA-B (B) HLA-C copy number prediction by LOH-HLA in PT, LR, B3 and B8.

Supplementary Figure 5. SCNA profile of plasma samples, B1-B2 and B4-B7

