## Supplementary figures and images for "Longitudinal whole-exome sequencing of cell-free DNA unravels the metastatic evolutionary dynamics of *BRCA2*-mutated breast cancer"

### Supplementary Figure 1a-b

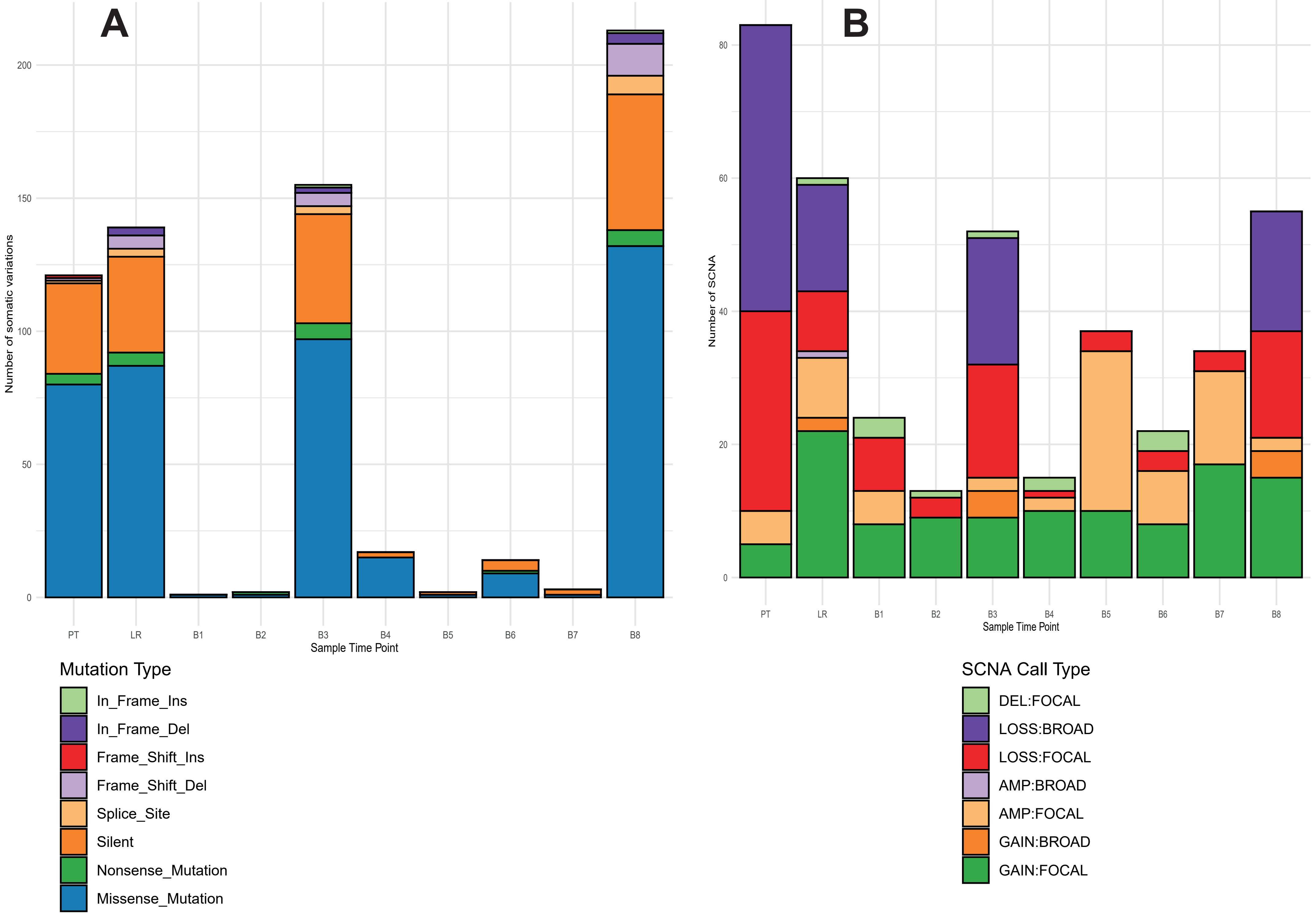

### Supplementary Figure 2

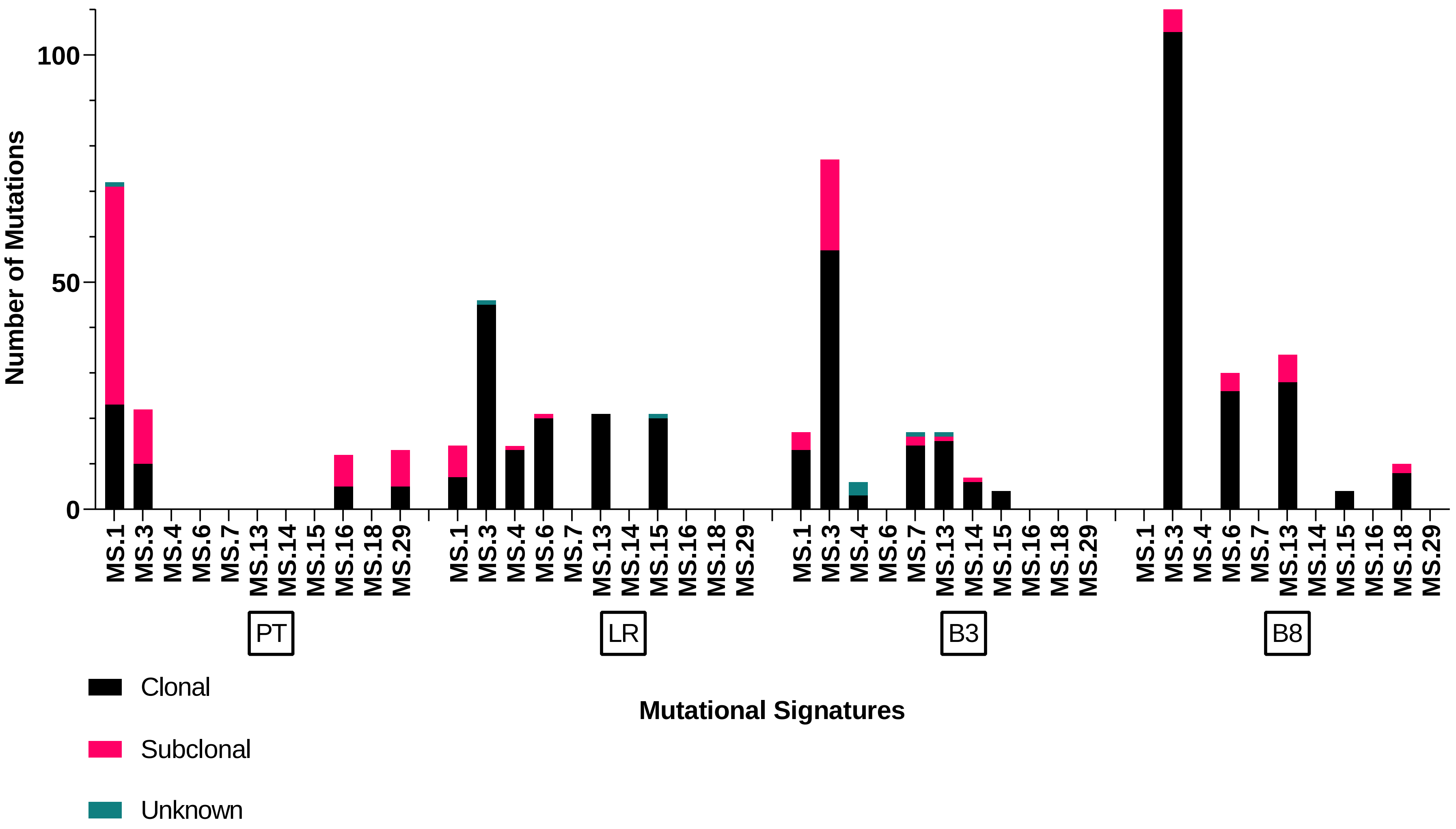

### Supplementary Figure 3a-b

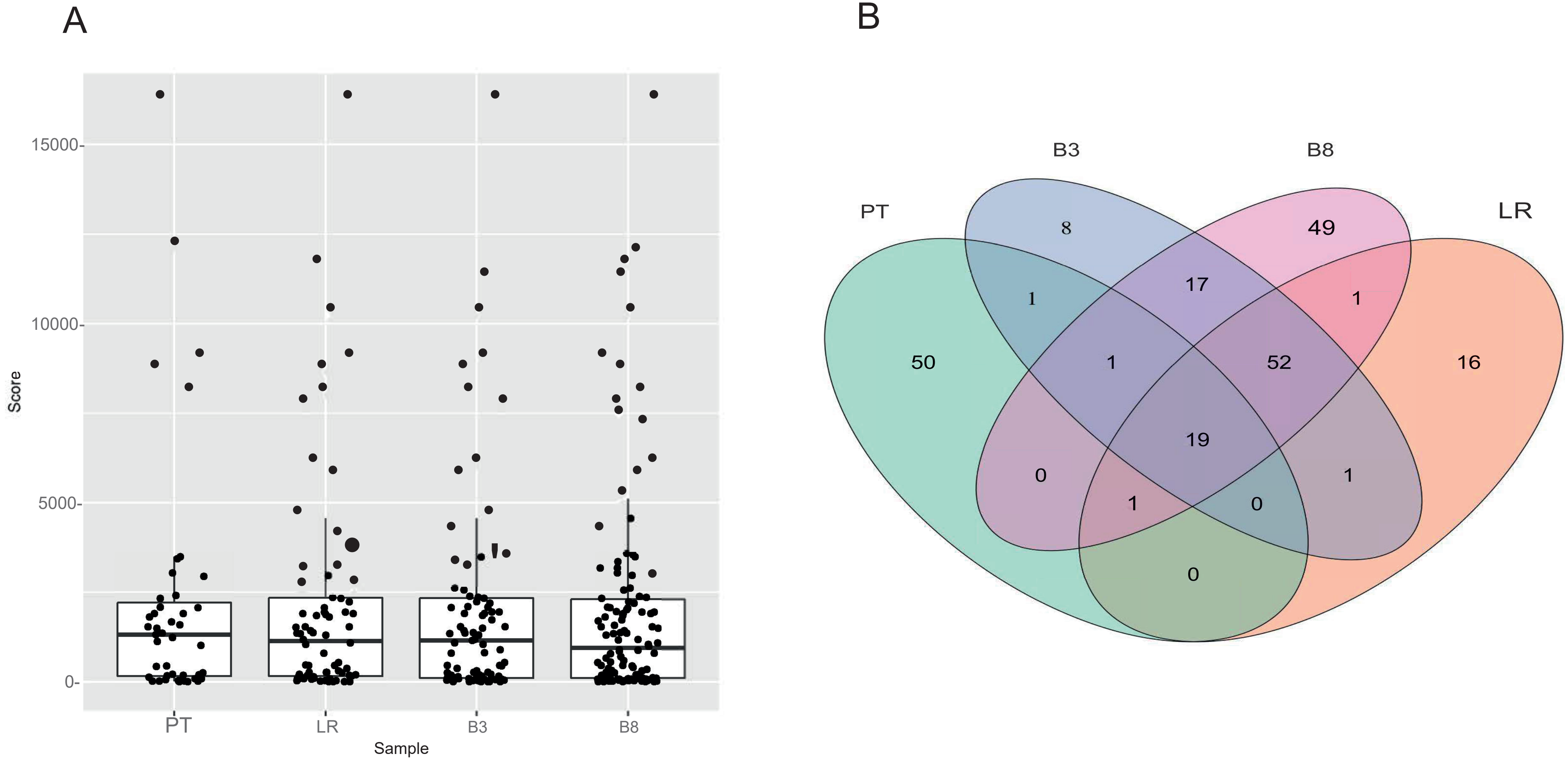

### Supplementary Figure 4

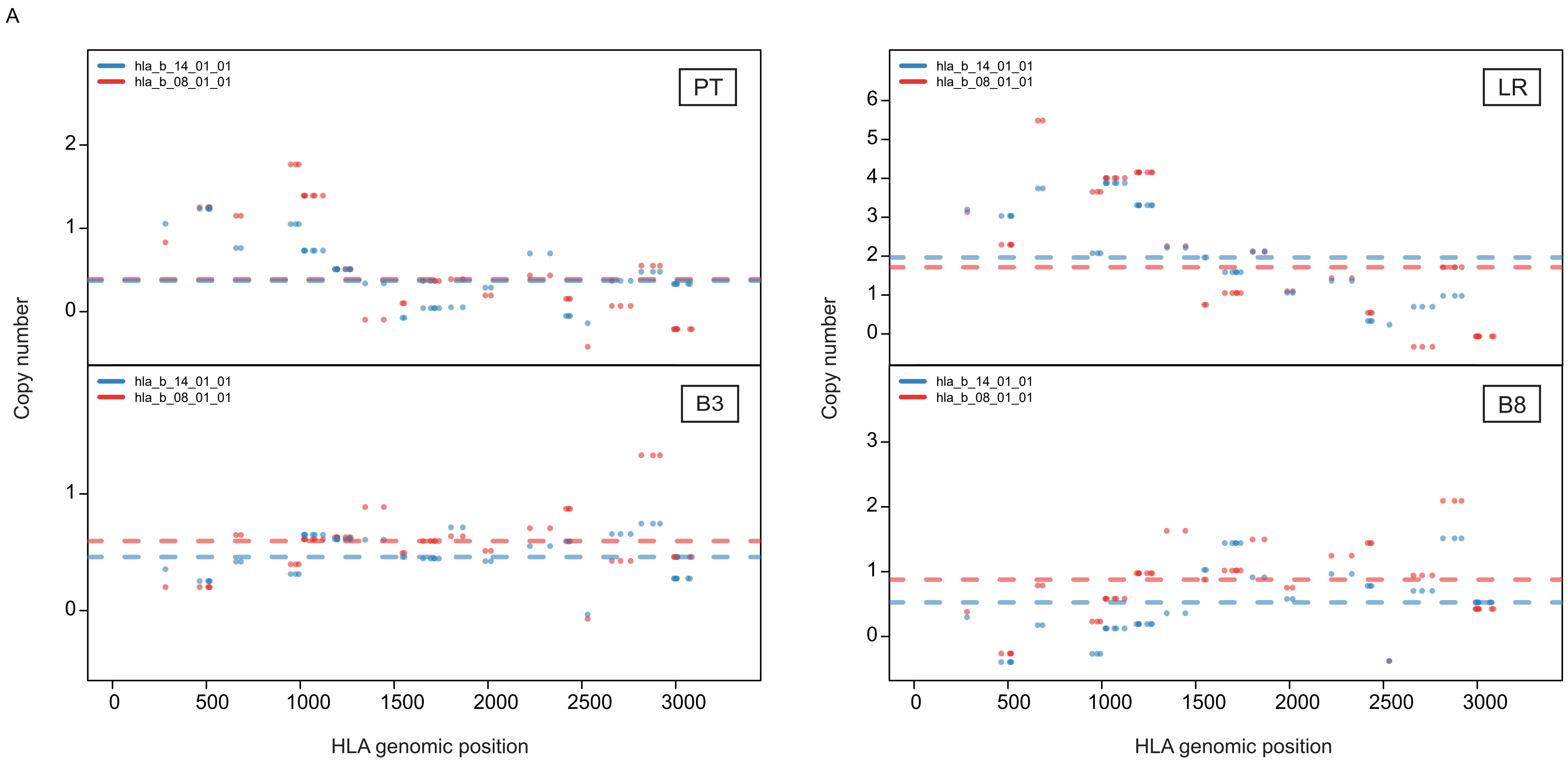

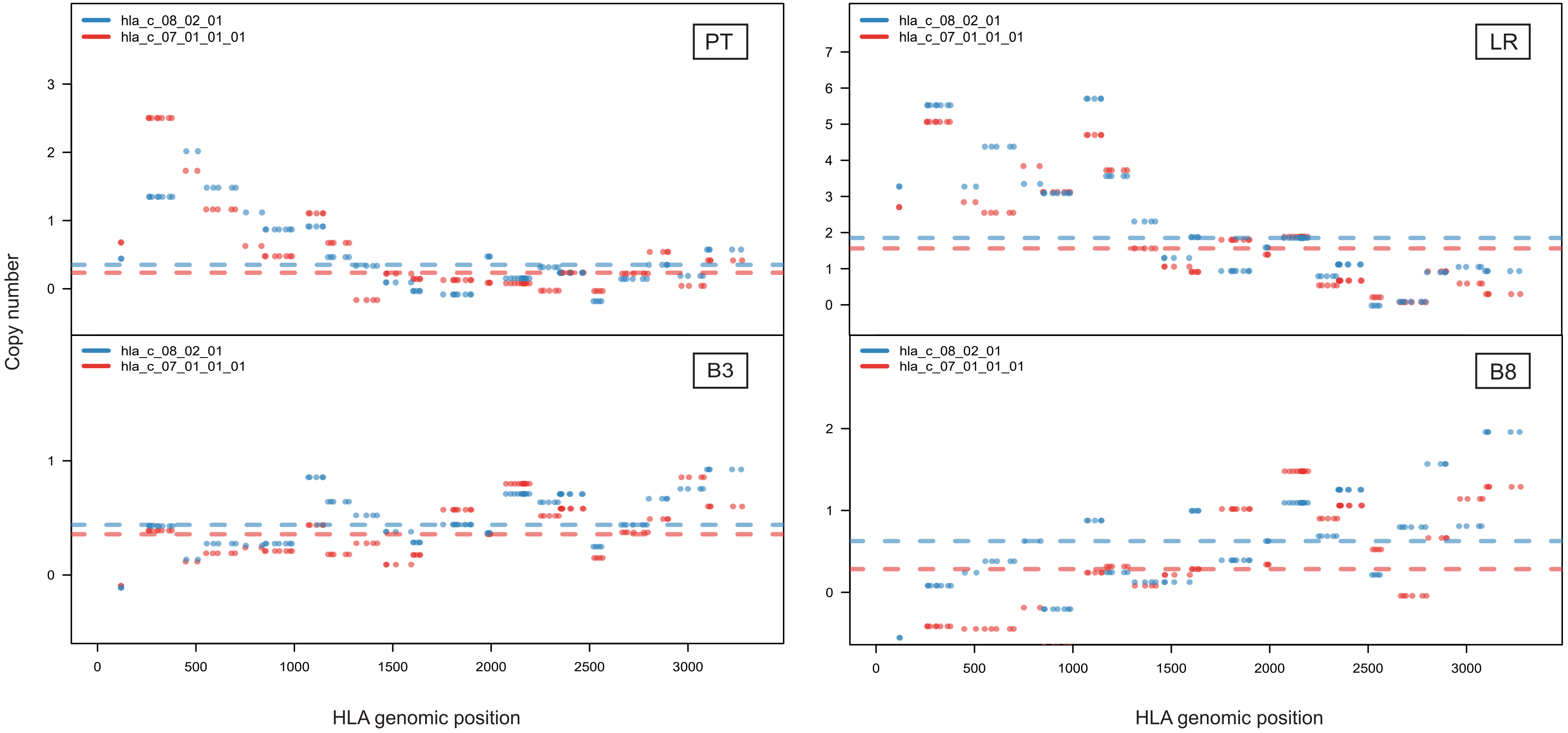

### Supplementary Figure 5

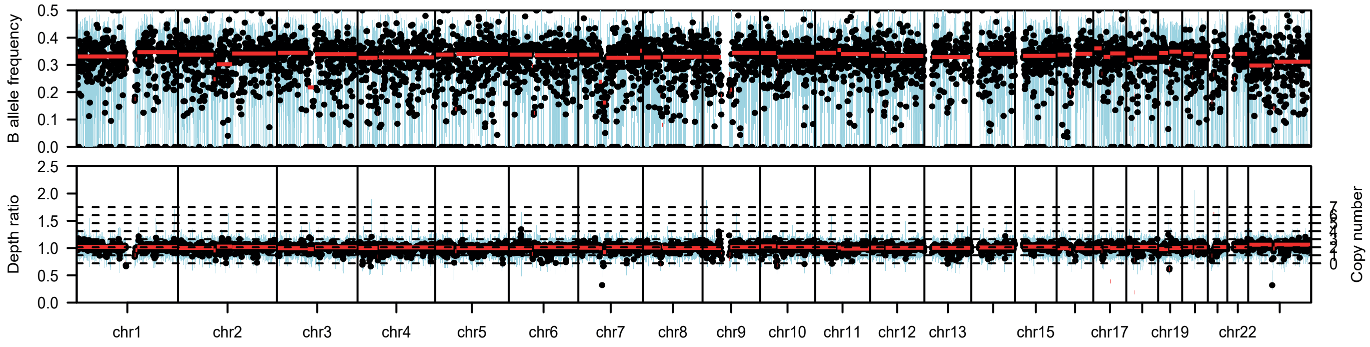

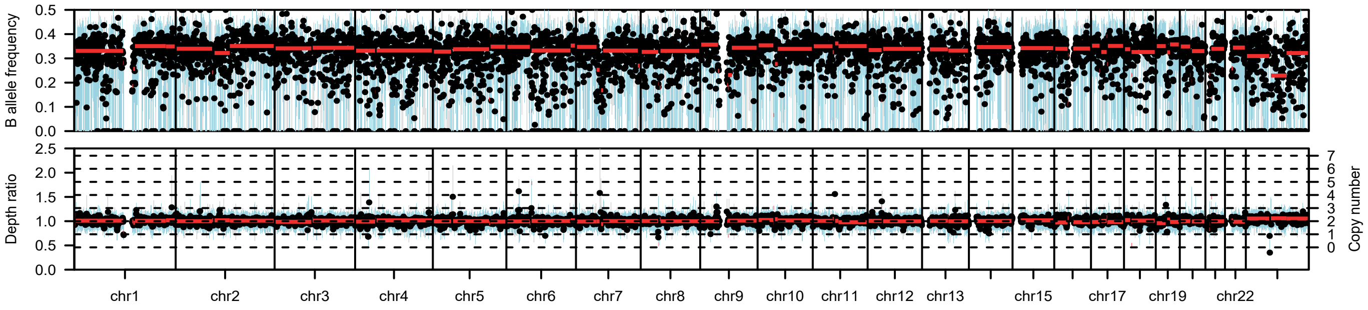

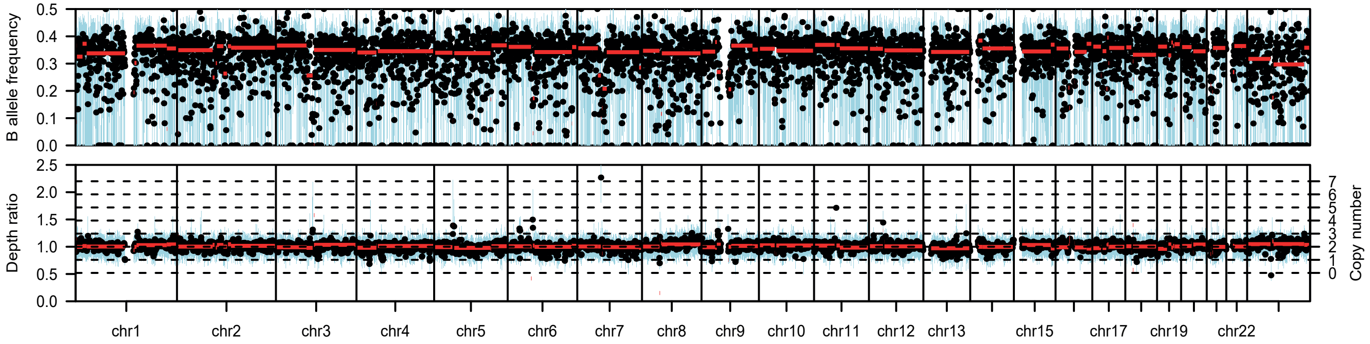

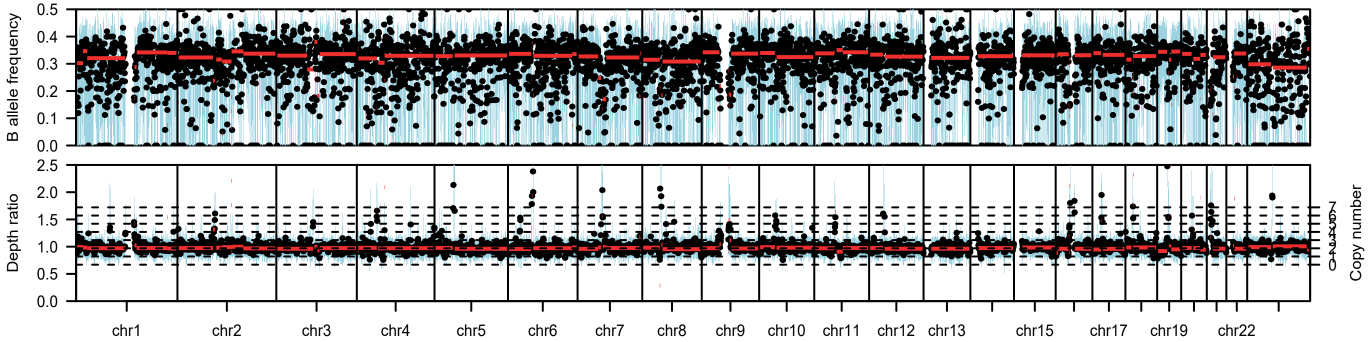

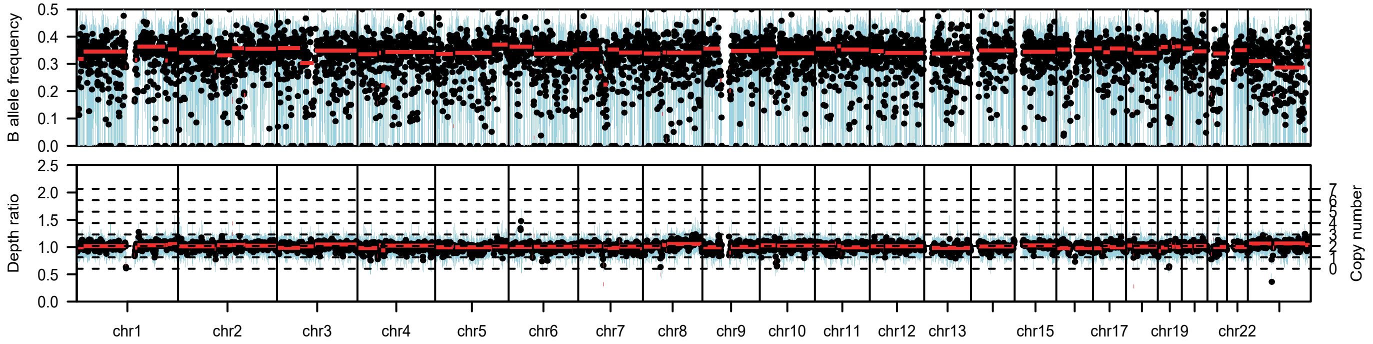

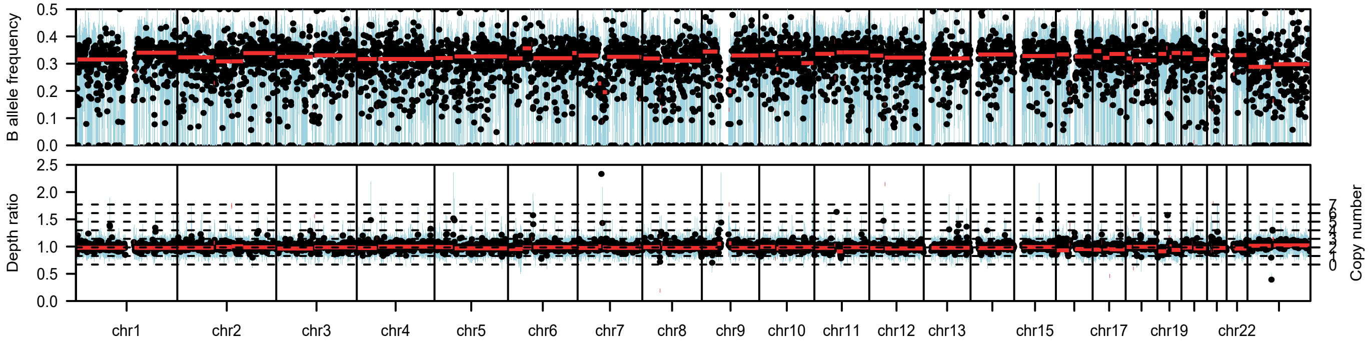
